# The genome sequence of the Lumpy Skin Disease virus from the outbreak in India suggests a distinct lineage of the virus

**DOI:** 10.1101/2022.09.15.508131

**Authors:** Lenin Bhatt, Rahul C. Bhoyar, Bani Jolly, Ravi Israni, Harie Vignesh, Vinod Scaria, Sridhar Sivasubbu

## Abstract

Although previously confined to regions within Africa, Lumpy Skin Disease Virus (LSDV) infections have caused significantly large outbreaks in several regions of the world in very recent years. In 2019, an outbreak of the disease was reported from India with low rates of morbidity and no reported mortality. However, in 2022, an ongoing outbreak of LSDV spanning over 7 states in India resulted in a reported mortality of over 80,000 cattle over a short period of three months. Here, we report complete genome sequences of 6 viral isolates of LSDV collected from affected cattle during an ongoing outbreak of the disease in Rajasthan, India. Analysis of the viral sequences suggests the genomes from the 2022 outbreak harbour a large number of genetic variations compared to the reference genome and form a distinct lineage. This report also highlights the importance of genome sequencing and surveillance of transboundary infectious agents and genomic characterisation of outbreaks.

## INTRODUCTION

Lumpy Skin Disease (LSD) is a virus disease of cattle, with significant morbidity and mortality. The disease is characterised by nodular skin lesions evident as lumps along with lymphadenopathy, with severe cases resulting in the death of the affected animal (Shumilova et al., 2022). The morbidity rate in LSD varies widely from 50–100% (Tuppurainen and Oura, 2012). While in general, the mortality rate is usually low (less than 5%) much higher mortality rates have been reported (Molla et al., 2017). The disease was largely confined to Africa until 1981 with significant outbreaks in Asia and Eurasia in recent years. The disease is therefore categorised as a notifiable emerging transboundary disease by the World Organisation for Animal Health (OIE) (Sameea Yousefi et al., 2017). Recent years have seen a significant spread of the disease across the Middle East, Europe and Asia (Kumar et al., 2021). More recently in 2019, it has been reported in the Asia and Pacific region including India, China, Nepal, Sri Lanka, Bhutan and a large outbreak in Bangladesh (Azeem et al., 2022).

LSD is caused by the Lumpy Skin Disease Virus (LSDV), a poxvirus of the genus capripoxvirus. The genome sequence of the Neethling strain of the virus suggested that similar to other poxviruses, it has a ^151 kbp size genome encoding 156 open reading frames (Tulman et al., 2001). The virus has a limited host range and largely affects cattle (Shen et al., 2011). Large variability in the morbidity of the disease has been reported, with morbidity rates across different studies ranging from 3%-85% (Babiuk et al., 2008; Şevik and Doğan, 2017). The mortality rate for previous outbreaks of the disease has been observed to be low (1%-3%), although in some regions high mortality rates (over 40%) have also been reported (Davies, 1991; Irons et al., 2005; Babiuk et al., 2008). Mortality rates of the disease from more recent outbreaks in Iraq and Egypt were also reported to be low (<1%) (Al-Salihi and Hassan, 2015; Tasioudi et al., 2016).

In mid-2022, the states of Gujarat and Rajasthan in India saw a large outbreak of LSD which spread across over seven states in the country and reported mortality of over 80,000 cattle over a very short span of three months. During the 2019 outbreak of the disease in India, morbidity rates were reported to be as low as 7.1%, and no mortality was reported (Sudhakar et al., 2020).

As India is home to over 190 million cattle, one of the largest in the world (Department of Animal Husbandry & Dairying, 2019), controlling LSDV infections is therefore of significant importance to prevent the socio-economic impacts of the disease in the country. Here, we report complete genome sequences of LSDV from the current outbreak of the disease, isolated from the affected cattle in India.

## METHODS

Samples were collected from five affected animals, which had symptoms suggestive of Lumpy Skin Disease during an ongoing outbreak of the disease in Rajasthan. Nasal swabs or skin scabs or both were collected by trained veterinary professionals. DNA extraction from the collected specimens was performed in biosafety level 2 laboratory standards using the automated nucleic acid extraction protocol (Genolution, Korea). Sequencing-ready libraries were constructed by adopting the Illumina tagmentation technique with minor modifications to the original protocol (Bhoyar et al., 2021). After quality checks, the synthesised libraries were sequenced on the NovaSeq 6000 platform (Illumina, USA) with 150X2 base-paired end reads.

The raw FASTQ files for each sample generated by the sequencer were trimmed using Trimmomatic (v0.39) to remove adapters and bases below the quality score of Q30 (Bolger et al., 2014). The quality controlled reads for individual samples were aligned to the Lumpy skin disease virus NI-2490 genome (NC_003027.1) using hisat2 (v2.1) to generate a reference-based mapping for the isolates (Tulman et al., 2001; Kim et al., 2015). Variant calling and consensus sequence generation were performed using VarScan (v2.4.4) and seqtk (v1.3) respectively (Koboldt et al., 2009). Only variants having a variant allele frequency by read count greater than 50% were considered. Variant calls for all individual samples were annotated using a custom database file created using ANNOVAR (Wang et al., 2010). For comparative analysis, 57 LSDV FASTA sequences deposited in GenBank with nucleotide completeness >90% were downloaded (Sayers et al., 2019). Variant calling and annotation for the downloaded sequences were performed using snp-sites and ANNOVAR respectively.

Phylogenetic analysis of the samples was done using the 57 genomes from GenBank and the genome NC_003027.1 as the root. A multiple sequence alignment file was generated using MAFFT (v7.505) and the phylogenetic tree was constructed using iqtree (v2.2.0.3) (Katoh and Toh, 2008), (Nguyen et al., 2015).

## RESULTS

A total of 8 viral isolates collected from 5 affected animals were processed for genome sequencing. Genome assemblies having a high percentage of genome coverage (>99%) were considered for further analysis, which resulted in 6 genome isolates. The isolates were assembled at an average genome coverage of 99.96% with a mean coverage depth of 596.39X (**Supplementary Data**).

A total of 177 unique variants were detected across the 6 isolates, of which 86 were synonymous single nucleotide variants (SNVs), 58 were nonsynonymous, 27 were indels, and 1 was a stopgain SNV while 5 variants were found in untranslated regions (UTR). A maximum number of variants were detected in the gene LSDVgp134 (N=7) (**Supplementary Data**). On comparison with 57 LSDV genome sequences downloaded from GenBank, we found that 47 variants detected in the 6 sequenced isolates were novel and were not present in any other genome sequence globally (**Figure 1A**). Of the novel variants, 14 were nonsynonymous SNVs while 27 are indels. A maximum number of variants were detected in the gene LSDVgp134 (N=3) and LSDVgp011 (N=3) while the most number of indels were found across 4 genes, LSDVgp026, LSDVgp067, LSDVgp134 and LSDVgp151 all of which had two indels each (**Supplementary Data**). None of the variants found in 4 genome sequences from India belonging to the 2019 outbreak of the disease deposited in GenBank was present in the 6 samples from the current outbreak (Kumar et al., 2021).

**Figure 1.**
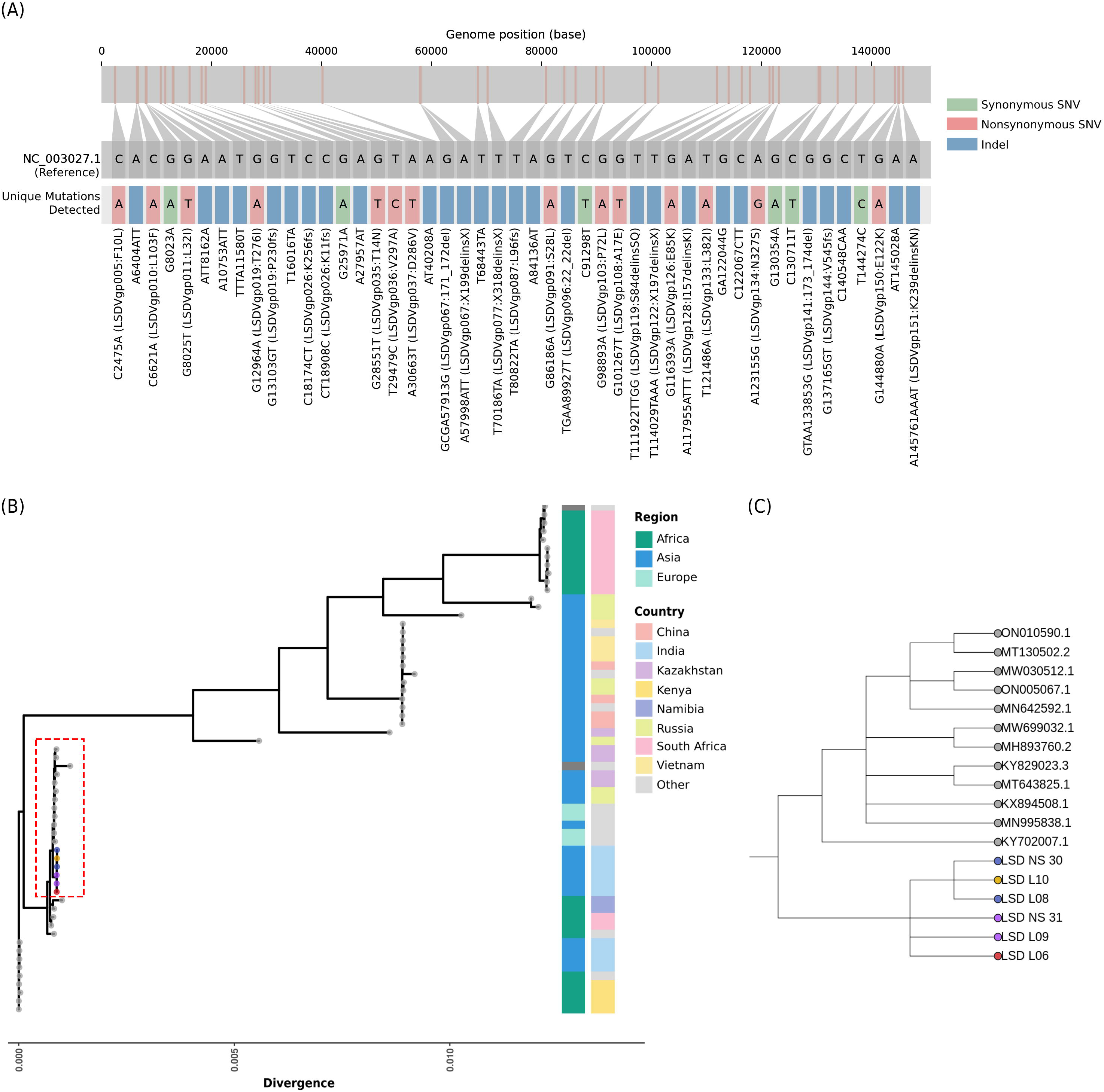
**(A)** Novel variants found across 6 genome isolates of LSDV as compared to global genomes deposited in GenBank. **(B)** Phylogenetic context of the 6 viral isolates of LSDV. Tree nodes belonging to the same host animal have been shown in the same colour. **(C)** The closest cluster of genomes (highlighted in **1B**) for the 6 viral isolates on the phylogenetic tree

Phylogenetic analysis showed that the 6 genome isolates from the present outbreak form a distinct cluster on the phylogenetic tree with little similarity to other global genomes (**Supplementary Data**). The cluster is defined by 39 unique variants which were present in all 6 isolates but not present in any other global genome. The phylogenetic context of the samples has been summarised in **Figure 1B**. The closest cluster for the 6 isolates on the phylogenetic tree comprises 12 sequences deposited from 7 Asian and European countries with collection dates ranging from 2012 to 2022 (**Figure 1C**).

Genome sequences of viruses isolated from both nasal swabs and skin scabs were available for 2 of the 5 host animals, which allowed us to check for intra-host variations. Analysis of the variants suggested that one nonsynonymous variant (A30663T LSDVgp037:D286V) which was present in the viral isolated from skin scab for 1 animal which was not present in the nasal swab isolate of the same host sample potentially pointing to intra-host evolution. The variant position had a good depth of coverage (>100X) for both samples.

## DISCUSSION AND CONCLUSIONS

Being the largest producer of milk in the world, the spread of LSD across the cattle population in India thus has devastating effects on the agrarian economy as well as the livelihoods of dairy farmers (FAO, UN). Although other capripoxvirus infections including sheep pox and goat pox are endemic in India, LSDV infections were considered to be constrained to the Sub-Saharan African region till 1981 (Das et al., 2021). As the virus continues to spread and evolve, genomic characterization of LSDV is thus useful for understanding the epidemiology and evolution of the virus.

In this study, we have reported the whole-genome sequences of 6 viral isolates of LSDV that were collected from the state of Rajasthan during an ongoing outbreak of infections in India. Analysis of the 6 viral isolates shows that the genomes from the 2022 outbreak of the disease form contain a large number of genetic variants as compared to previous genomes available in the public domain. The presence of an additional variant (LSDVgp037:D286V) in the virus isolated from skin scab as compared to nasal swab of the one host animal is suggestive of potential intra-host evolution of LSDV.

The genome sequences also form a distinct cluster on the phylogenetic tree of all publicly available LDSV genome sequences and bear limited similarity to other global genomes. Due to the limited number of genome sequences available for LSDV, the source of the outbreak could not be traced, further suggesting that additional genomes for the virus could help uncover potential outbreaks and connect existing outbreaks that are apparently unrelated.

Put together, the study iterates that genomic surveillance for characterising circulating strains of important transboundary infectious agents during ongoing outbreaks, such as LSD, is essential for early detection of the disease as well as for formulating interventions for disease control.

## ACKNOWLEDGEMENTS

Authors acknowledge the funding support from project codes CLP0040 and CNP0007. BJ acknowledges a research fellowship from CSIR India. The authors acknowledge investigators for the data deposited in the public domain. A detailed list of acknowledgements is available in the Supplementary Data file.

## AUTHOR CONTRIBUTIONS

RI and LB examined the animals, performed the clinical evaluation and collected the samples. RB and HV performed the library preparation and sequencing. BJ performed the data analysis. VS and SSB conceptualised the project and provided oversight. All authors contributed to writing the manuscript.

## DATA AVAILABILITY

The data that supports the findings of this study is available in the NCBI Short Read Archive (BioProject ID PRJNA880745). The FASTA sequences for the isolates reported in the study are available at https://github.com/banijolly/LSDV_India.git.

